# Chloroplast transit peptides often require downstream unstructured sequence in *Chlamydomonas reinhardtii*

**DOI:** 10.1101/2021.11.26.470094

**Authors:** OD Caspari

## Abstract

The N-terminal sequence stretch that defines subcellular targeting for most nuclear encoded chloroplast proteins is usually considered identical to the sequence that is cleaved upon import. Yet here this study shows that for nine out of ten tested Chlamydomonas chloroplast transit peptides, additional sequence past the cleavage site is required to enable chloroplast targeting. Using replacements of native post-cleavage residues with alternative sequences points to a role for unstructured sequence at mature protein N-termini.

## 2 Introduction

Chloroplasts derive from once free-living cyanobacteria and retain genomes of their own, yet the vast majority of genes coding for plastid proteins are located in the nucleus (Shi and Theg, 2013). Proteins are expressed in the cytosol as preproteins containing N-terminal chloroplast targeting peptides (cTP), which direct import into the chloroplast (Chotewutmontri et al., 2017). Upon import, the N-terminus of the preprotein is cut off at a defined cleavage site and degraded (Teixeira and Glaser, 2013; Kmiec et al., 2014).

The cTP, i.e. the sequence both necessary and sufficient to generate chloroplast targeting, is generally assumed to coincide with the sequence stretch that is cleaved off. It is this cleaved presequence that has been studied for decades, with a well-documented tripartite structure and multiple sequence elements now recognized to contribute to targeting (von Heijne et al., 1989; Chotewutmontri et al., 2017; Caspari and Lafontaine, 2021). Yet the cleavage site may not always be a reliable guide for defining the cTP. Parts of the mature protein N-terminus have been recognized before as a requirement for efficient targeting in higher plants (Bionda et al., 2010; Rolland et al., 2016; Shen et al., 2017). Bionda et al. (2010) described a length requirement of ~55-60 amino acids for *Arabidopsis thaliana*, and proposed that in the ~50% of chloroplast targeted proteins where the cleavable part of the cTP is shorter, residues C-terminal of the cleavage site contribute unstructured sequence towards targeting. There are even examples of import through the canonical TOC/TIC system of proteins lacking a cleavable N-terminal presequence altogether (Chang et al., 2014), implying the entire targeting information is present in the mature protein.

It was noted decades ago that cleavable cTPs are generally shorter in the model green alga *Chlamydomonas reinhardtii* compared to higher plants based on the placement of cleavage sites (Franzén et al., 1990), suggesting there may be a role for mature protein N-termini. In Chlamydomonas, the Rubisco activase (RBCA) cTP has recently been found to require post-cleavage site residues to target a fluorescent reporter protein to the chloroplast (Garrido et al., 2020), in line with earlier work regarding Rubisco small subunit (RBCS) cTP from Chlamydomonas (Razzak et al., 2017) and higher plants (Comai et al., 1988). This present study revisits the role of post-cleavage site residues, and finds that substantial stretches of unstructured sequence appear to be required to enable targeting by most Chlamydomonas cTPs.

## 3 Materials and Methods

Constructs were Gibson assembled as described in (Garrido et al., 2020) based on pMO611, a variant of published pMO449 (Onishi and Pringle, 2016) where an upstream start codon was mutated to CTG. Sequences inserted upstream of Venus were amplified from Chlamydomonas genomic DNA extracted from wild-type strain T222+, with the following exceptions: construct ‘RBCS-cTP+23’ was amplified from cDNA generated from T222+ to avoid introducing an intron not present in the other RBCS-cTP constructs; the G-rich artificial linker sequence was amplified from plasmid CrRaf1-pJHL encoding Raf1-strep (Wietrzynski et al., 2021). Corresponding amino acid sequences are given in Table 1. Constructs were verified by sequencing (Eurofins). Transformants were obtained and microscopy performed as described before (Garrido et al., 2020).

**Table 1.**
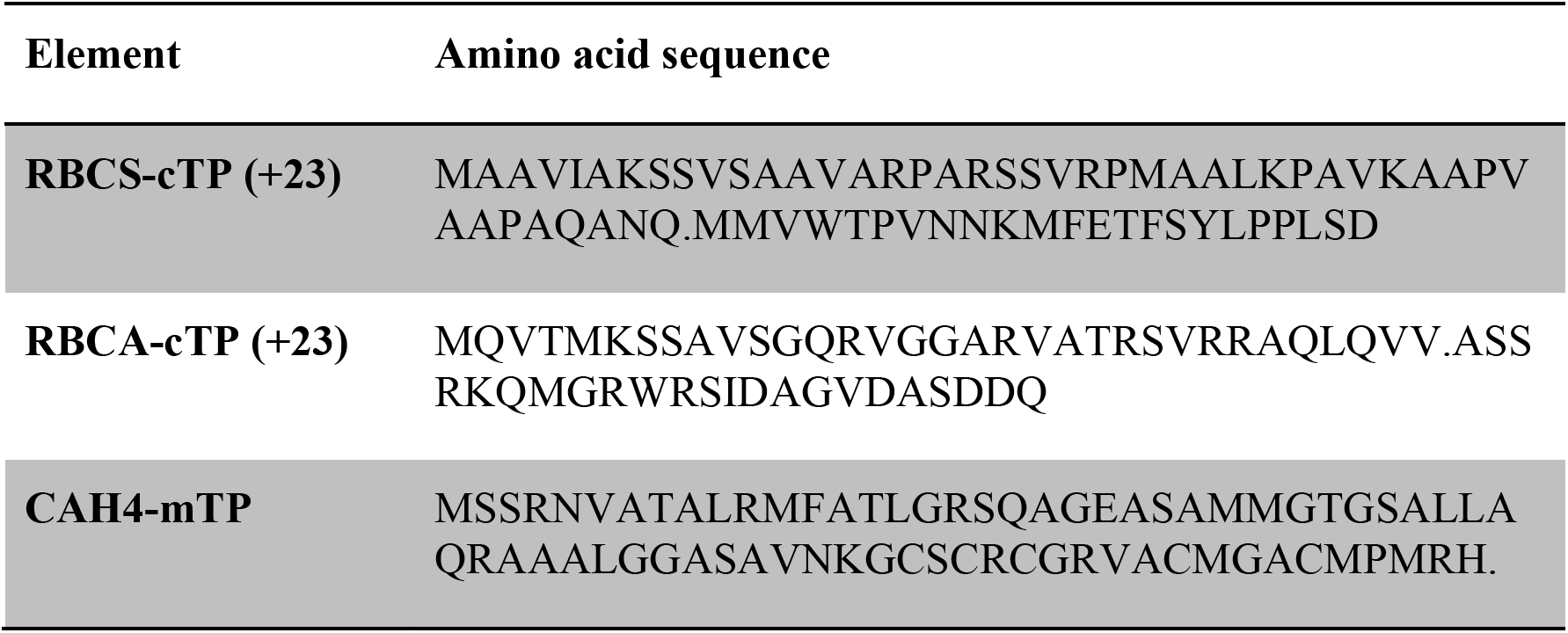
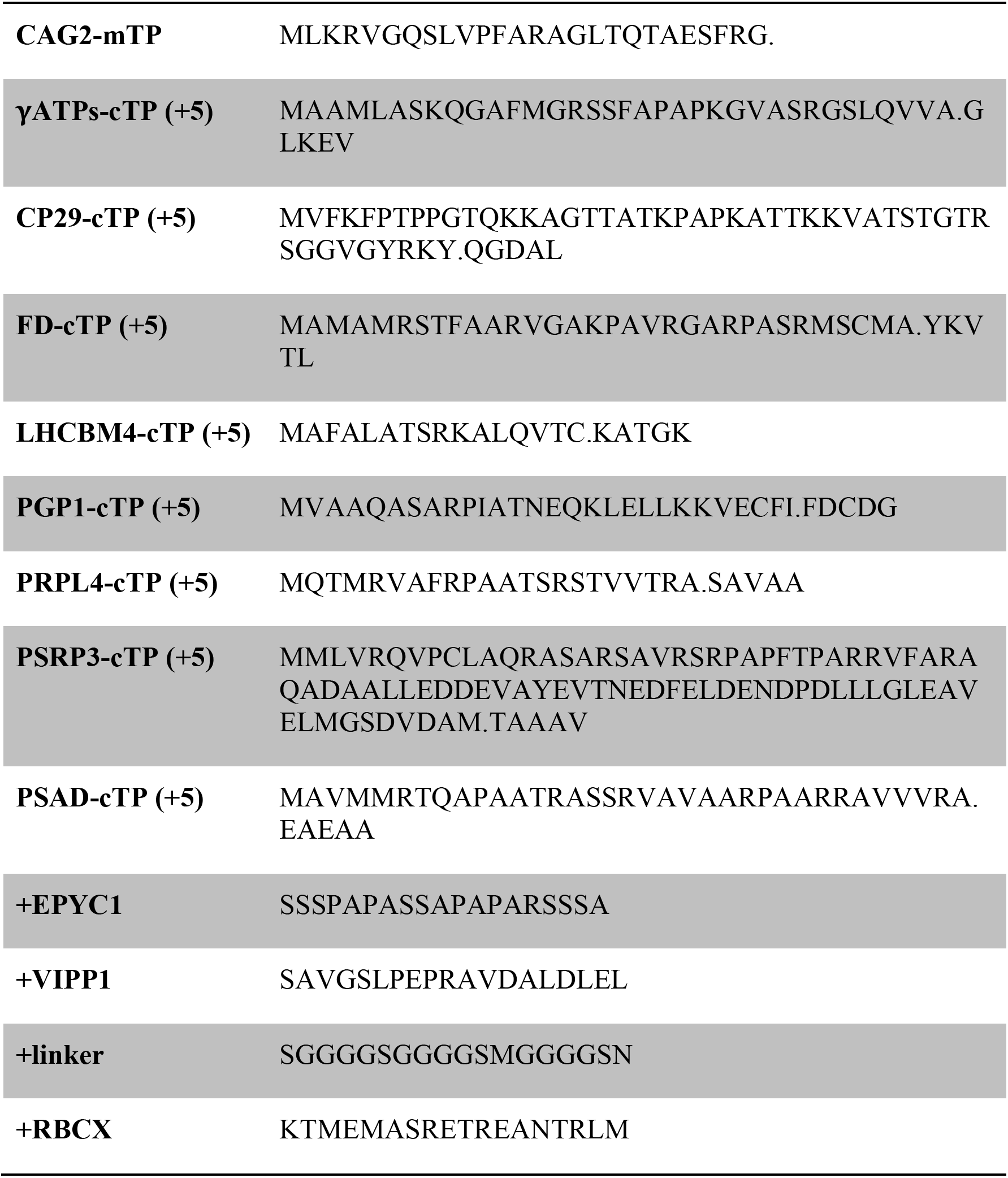
Amino acid sequences. Cleavage sites are indicated with a full stop between residues.

## 4 Results

Native cTPs were tested by inserting the respective genomic sequences upstream of a Venus fluorescent reporter in a bicistronic expression vector (Onishi and Pringle, 2016; Caspari, 2020). Neither RBCA nor RBCS cTPs enabled import into the chloroplast (Fig. 1A,B), instead showing Venus fluorescence from across the cytosol or from within globular structures outside the chloroplast respectively. Given that successful targeting using the RBCS-cTP including 23 residues downstream of the cleavage site had previously been reported in Chlamydomonas (Razzak et al., 2017) and higher plants (Comai et al., 1988) and extending the RBCA-cTP by 23 residues had equally restored targeting (Garrido et al., 2020), a series of successive extensions by 3 (Fig. 1C,D), 5 (Fig. 1E,F), 10 (Fig. 1G,H) and 23 residues (Fig. 1I,J) was prepared. The +3, +5 and +10 constructs of both RBCA-cTP and RBCS-cTP all show significant Venus fluorescence from outside the chloroplast (Fig. 1C-H), and only the RBCA-cTP+10 construct also shows some partial localization of Venus to the chloroplast (Fig. 1G). By contrast, the +23 constructs show clear accumulation of Venus inside the chloroplast (Fig. 1I,J), strikingly visible as fluorescence from inside the pyrenoid, a proteinaceous structure within the chloroplast that generates a visible dip in chlorophyll fluorescence (Mackinder et al., 2017). Note that the RBCA-cTP+23 construct contains a pyrenoid-localisation motif (Meyer et al., 2020). By contrast, mitochondrial targeting peptides (mTPs) from carbonic anhydrase 4 (CAH4) and γ-carbonic anhydrase 2 (CAG2) show mitochondrial targeting, evident as co-localisation of Venus and MitoTracker fluorescence, in the absence of any post-cleavage site residues (Fig. 1K,L). CAG2-mTP shows some remaining signal from the cytoplasm, which persists when the construct is extended by 23 residues past the cleavage site (Garrido et al., 2020).

**Fig. 1.**
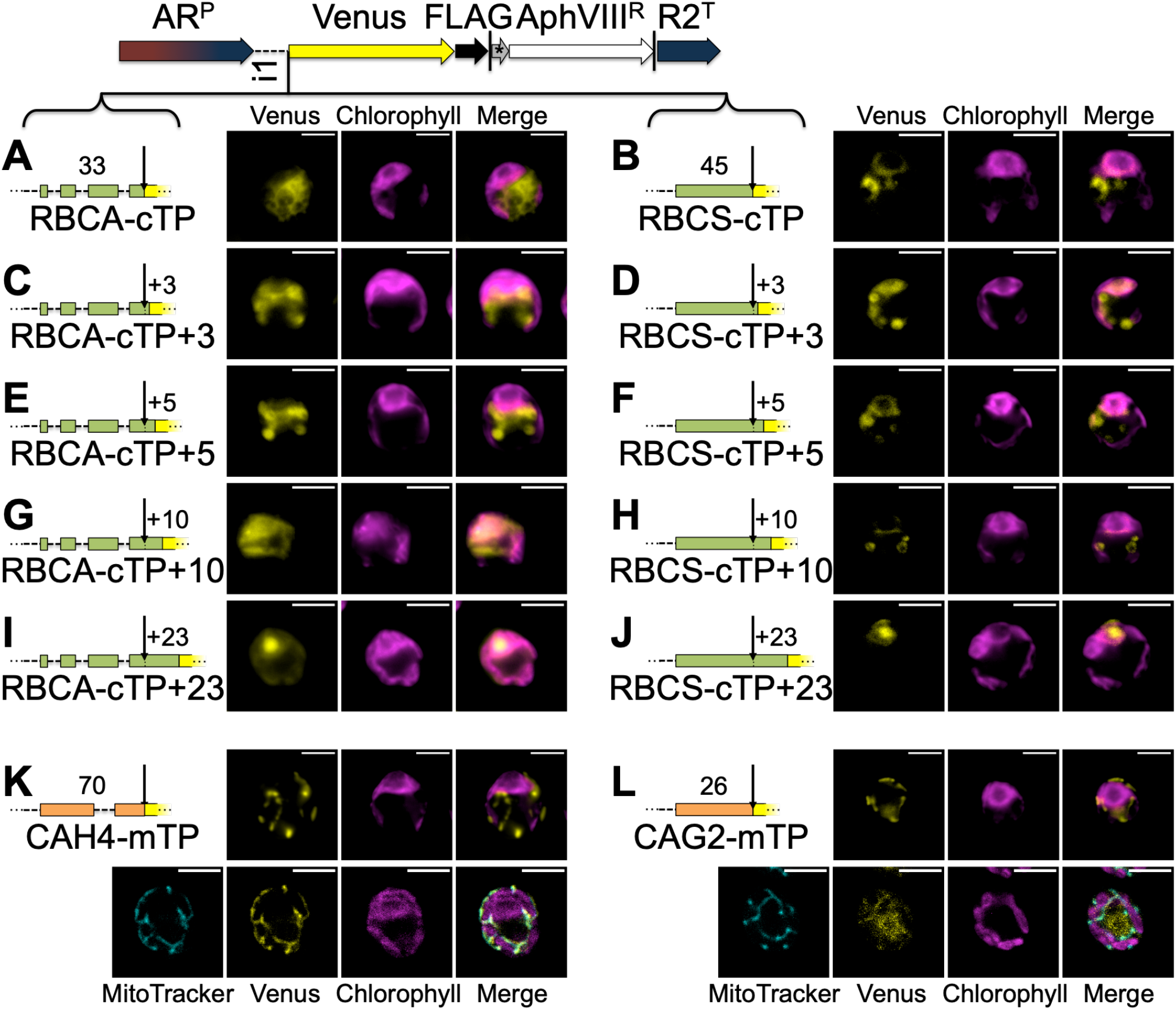
A stretch of post-cleavage site residues is necessary for chloroplast targeting peptides. Constructs were generated by inserting candidate peptides upstream of Venus in a bicistronic expression vector made of the following components: AR^P^: Hybrid *HSP70a-RBCS2* promoter, i1: *RBCS2* intron 1, Venus: Venus fluorescent protein (a YFP variant), FLAG: triple-FLAG tag, |: stop codon, *: bicistronic intergenic sequence (tagcat), AphVIII^R^: Paromomycin resistance gene, R2^T^: *RBCS2* terminator. Candidate peptides are Rubisco activase (RBCA, A-E) and Rubisco small subunit (RBCS, B-J) chloroplast targeting peptides (cTP) extended by 0, 3, 5, 10 or 23 residues beyond the cleavage site into the mature protein, as well as carbonic anhydrase 4 (CAH4, K) and γ-carbonic anhydrase 2 (CAG2, L) mitochondrial targeting peptides (mTP). Lengths in amino acid residues are indicated above each peptide. Introns within peptides are symbolized by dotted lines. Typical epifluorescence images of resulting transformants in strain T222^+^ show fluorescence from the Venus channel in yellow and chlorophyll autofluorescence in magenta. Confocal microscopy images include an additional MitoTracker channel in cyan for mTPs. All scale bars are 5μm in length.

To test whether the presence of a relatively long stretch of mature protein is a general requirement rather than an exception concerning specifically RBCA and RBCS, a further eight cTPs were assayed for import (Fig. 2). These cTPs were each extended by 5 residues beyond the cleavage site to ensure any lack of import was not an artifact arising from a missing cleavage site, for which several residues past the cleavage site seem to be important (Tardif et al., 2012). Only one of the tested cTP+5 constructs, that of PSAD, generates chloroplast localized Venus signal (Fig. 2A-H). Chloroplast targeting with the PSAD-cTP also works in the absence of any post-cleavage site residues (Fig. 2I).

**Fig. 2.**
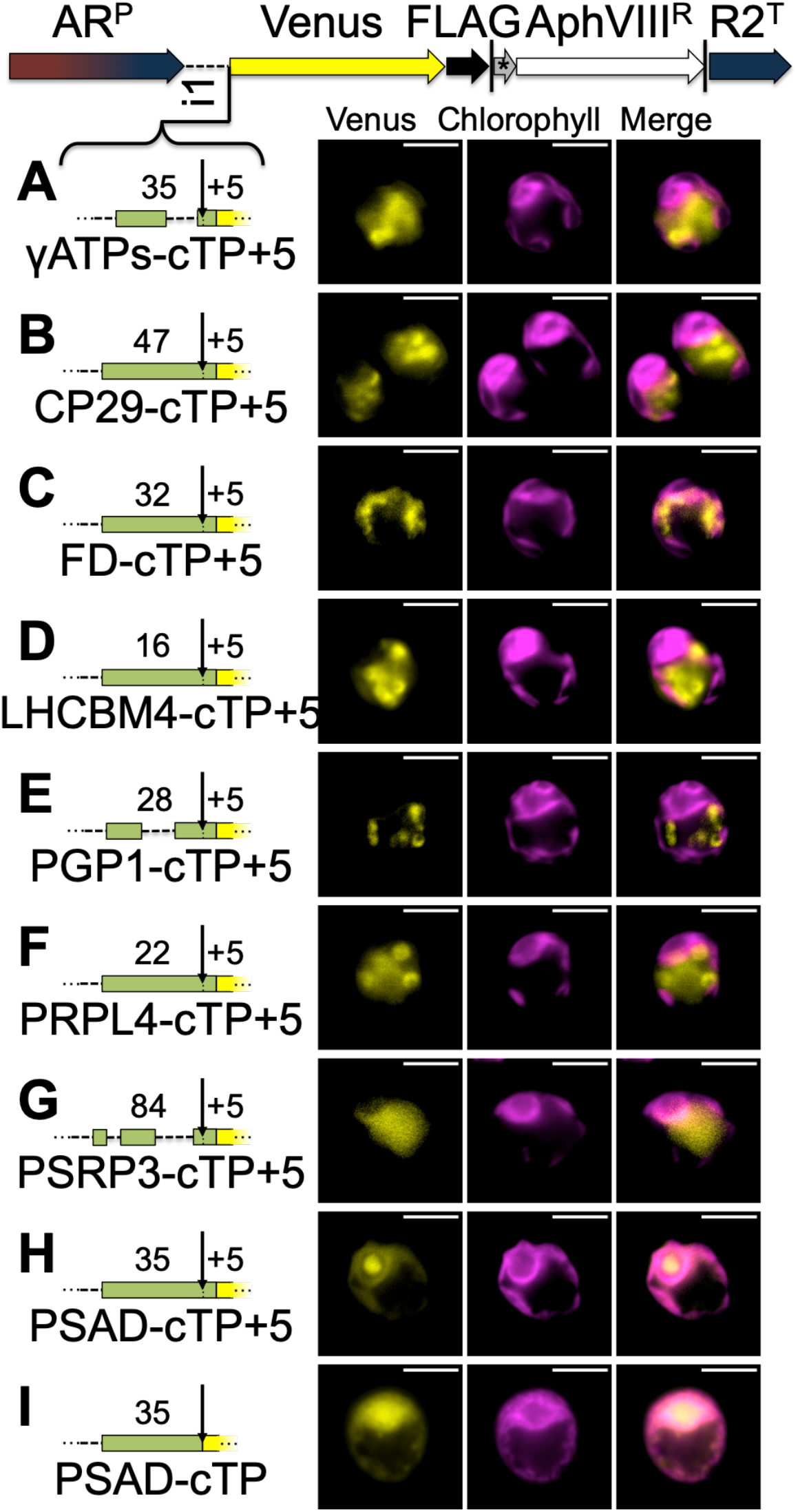
Without a stretch of post-cleavage site residues, Chlamydomonas chloroplast targeting peptides fail to import. Candidate chloroplast targeting peptides, each including 5 post-cleavage site residues, from (A) chloroplast ATP-synthase gamma chain (γATPs), (B) chlorophyll a/b binding protein of photosystem II (CP29), (C) ferredoxin (FD), (D) chlorophyll a/b binding protein of light harvesting complex II (LHCBM4), (E) phosphoglycolate phosphatase (PGP1), (F) plastid ribosomal protein L4 (PRPL4), (G) plastid-specific ribosomal protein 3 (PSRP3) and (H) Photosystem 1 subunit D (PSAD), were inserted in-frame upstream of Venus. (I) A second PSAD construct lacking post-cleavage site residues was probed. Lengths in amino acid residues are indicated above each peptide. Representation as in Fig. 1.

To follow up the hypothesis that post-cleavage site residues contribute unstructured sequence towards chloroplast targeting, four constructs were prepared that each extend the RBCA-cTP+5 by different sequences (Fig. 3). Five mature RBCA residues were kept to ensure the cleavage site would be maintained intact. Three constructs contained unstructured sequence downstream (A-C) and were expected to generate chloroplast targeting, whereas a fourth (D) contained a α-helical stretch that was expected to impede chloroplast import. Two of the unstructured sequences as well as the helical stretch were copied from Chlamydomonas nuclear genes encoding chloroplast-localized proteins EPYC1 (also known as LCI5), VIPP1 and RBCXa respectively. In each of these cases, sequence was chosen from the C-terminal half of the protein, to avoid any sequence that might contribute to the cTPs of these proteins. In addition, a standard G-rich linker sequence, such as is commonly used to join elements when generating fusion proteins in molecular biology, was used to test whether an artificial flexible sequence could substitute for a native unstructured one.

**Fig. 3.**
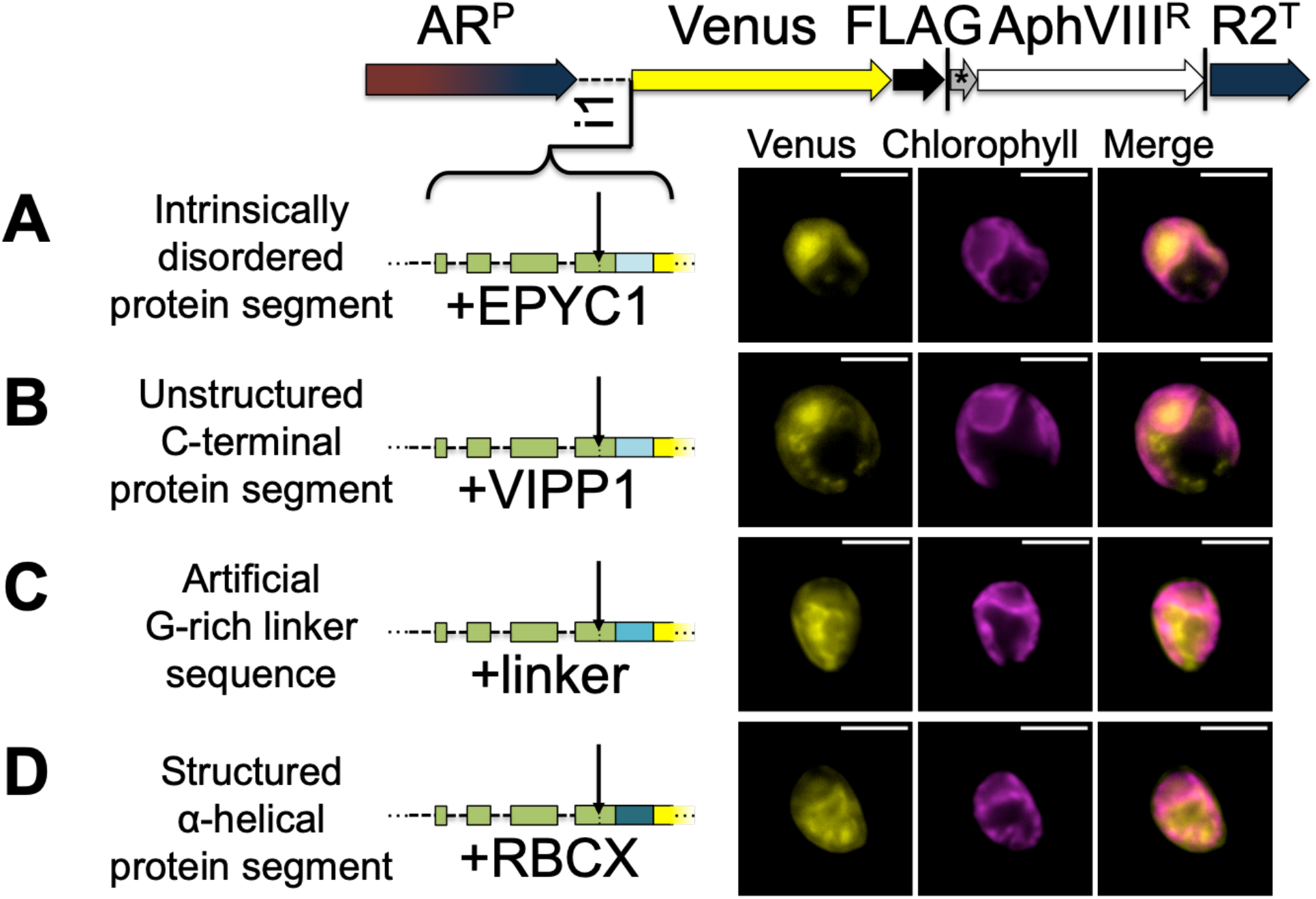
Unstructured downstream sequence aids cTP targeting. The following candidate sequences were inserted downstream of RBCA-cTP+5 and upstream of Venus: (A) residues 195-213 of intrinsically disordered protein EPYC1, which are part of the 3^rd^ out of 4 repeats; (B) residues 260-278 of VIPP1, which form part of an unstructured C-terminus; (C) an artificial linker sequence mainly consisting of quintuple-G interspersed with S; (D) residues 135-154 of RBCXa, which form part of the most C-terminal α-helix. Refer to Table 1 for amino acid sequences. Representation as in Fig. 1.

Of these, natively unstructured sequences from the third repeat of intrinsically disordered EPYC1 and the disordered C-terminus of VIPP1 generated chloroplast localization of the Venus reporter (Fig. 3A,B). Note that the VIPP1-derived sequence also shows additional Venus signal emanating from cytosolic structures (Fig. 3B). By contrast, both the artificial flexible linker and the C-terminal *α*-helix of RBCXa resulted in only a small amount of Venus localizing to the chloroplast, with a majority of signal emanating from structures outside the chloroplast (Fig. 3C,D).

## 5 Discussion

By definition, targeting peptides comprise the sequence that is both necessary for targeting a native cargo protein to its subcellular destination, and sufficient to redirect a heterologously expressed cargo to the same location (Chotewutmontri et al., 2017). The present study demonstrates that in Chlamydomonas, the amino-terminal sequence required to specify chloroplast targeting exceeds the cleavable part for nine out of ten tested cTPs. Care was taken to make the set of studied cTPs as representative as possible within the limited choice of cTPs with a well-characterized cleavage site (Terashima et al., 2011; Tardif et al., 2012). In particular, studied cTPs were chosen to include proteins related to photosynthesis (e.g. RBCA, RBCS, PSAD) as well as to plastid housekeeping (e.g. ribosomal proteins PRPL4, PSRP3), as import of these groups may be mediated by different receptors at the translocon (Chotewutmontri et al., 2017). The cleavage site should thus no longer be considered a reliable guide for identifying the N-terminal stretch containing all chloroplast targeting information.

The reduced length of algal cTPs (Franzén et al., 1990) may partly be accounted for by specific sequence elements required for targeting in plant cTPs being reduced in Chlamydomonas, such as a TOC-interacting FGLK motif (Chotewutmontri et al., 2017; Razzak et al., 2017; Caspari et al., 2021). Yet the requirement for post-cleavage site residues implies that the amino-terminal sequences contributing to targeting in Chlamydomonas are in fact longer than recognized to date, with sequence elements that play a role in import present past the cleavage site just as in short plant cTPs (Caspari et al., 2021). With lengths of 56 and 60 residues respectively, the import competent RBCA-cTP+23 and RBCS-cTP+23 constructs mirror the ~55-60 amino acid requirement Arabidopsis cTPs (Bionda et al., 2010). In this sense, algal cTPs may not be shorter than their plant counterparts after all.

Previous studies suggested that the contribution of mature protein N-termini towards targeting may simply be a supply of unstructured sequence (Bionda et al., 2010; Shen et al., 2017). Bionda et al. (2010) found that using tightly folded titin as cargo protein prevented chloroplast import by short cTPs unless first denatured. Shen et al., (2017) found that ~20 residues of unfolded sequence downstream of the cTP cleavage site were required to target two *E. coli* proteins into rice chloroplasts. While less tightly folded than titin, the Venus fluorescent reporter used as cargo protein in the present study contains a short, tightly folded α-helix at the N-terminus (Rekas et al., 2002), which may conceivably impair import in the absence of an unfolded spacer sequence. In principle, the chloroplast import machinery is capable of unfolding any protein for post-translational import, driven by an ATP-powered motor complex in the chloroplast stroma (Chotewutmontri et al., 2017). However, initiation of import requires the cTP to reach across both TOC and TIC passively before being able to contact stromal motor proteins. It is this initiation step that is most likely disrupted by the presence of structured elements too close to the N-terminus (Bionda et al., 2010). Mitochondrial import lacks similar constraints, as mitochondrial targeting peptides are guided across the outer membrane translocase using interactions with increasing affinity (Schatz, 1997; Fukasawa et al., 2017), and then make use of the proton motive force across the inner membrane to power import even before contacting the matrix-localized PAM complex that uses ATP to energize further import (Martin et al., 1991; Wiedemann and Pfanner, 2017). On the other hand, Razzak et al. (2017) report the presence of a specific motif (Q|MMVW) required for targeting in the Chlamydomonas RBCS-cTP that spans the cleavage site (Schmidt et al., 1979), suggesting post-cleavage site sequence elements may also play a more active role in specifying targeting.

Here, we found that chloroplast import by RBCA-cTP could indeed be restored using unstructured sequences taken from chloroplast proteins EPYC1 and VIPP1 as post-cleavage site spacers. EPYC1 is an intrinsically disordered protein comprised of four near-identical internal repeats that act as linker protein within the chloroplast pyrenoid (Mackinder et al., 2016), a proteinaceous structure within the chloroplast that is central to the algal carbon concentrating mechanism (Caspari et al., 2017). VIPP1 plays a role in the biogenesis of thylakoid membranes and has an unstructured C-terminus (Zhang et al., 2016; Theis et al., 2019). C-terminal sequences from these proteins are not expected to contain any specific targeting motifs, thus restoration of targeting lends further support to the importance of unstructured sequence at the N-terminus. In the same vain, copying an inherently structured sequence from the most C-terminal helix of RBCXa (Bracher et al., 2015) largely prevented chloroplast targeting (Fig. 3D).

An overall picture thus emerges of short algal cTPs up to the cleavage site containing the information for chloroplast targeting, but relying on additional unstructured sequence at mature protein N-termini to allow import to be successfully initiated. The importance of sufficient length and the presence of unstructured sequence elements for chloroplast targeting was equally evident in recent work aimed at retracing likely steps in the evolution of targeting peptides from putative antimicrobial origins (Caspari et al., 2021). Additional complexities likely exist, as several observations point to further factors being at play: A G-rich linker failed to restore chloroplast import beyond what had been seen for the RBCXa helix (Fig. 3C), suggesting that this artificial flexible sequence differs in important ways from native unstructured sequences. Furthermore, the very long PSRP3-cTP+5 construct (89 residues) failed to target whereas the relatively short PSAD-cTP (35 residues) achieved targeting, suggesting additional factors are in play beyond cTP length. The case of PSAD is particularly intriguing, as this cTP should be too short to reach into the stroma passively, yet appears to circumvent the impedance of a folded Venus cargo. Presumably, the cargo protein is kept in an unfolded state prior to import, e.g. through recruitment of chaperones interacting with the cTP, or perhaps through co-translational import following polysome recruitment to the chloroplast envelope by this cTP. Either way, for the purposes of sending heterologous proteins to the chloroplast (Crozet et al., 2018), the PSAD-cTP is likely the best choice among Chlamydomonas cTPs studied so far to ensure targeting without a need for additional sequence at the mature protein N-terminus.

## 7 Data availability statement

The original contributions presented in the study are included in the article/supplementary material, further inquiries can be directed to the corresponding author/s.

## 8 Author Contributions

Conceptualisation, Methodology, Investigation, Analysis, Interpretation, Visualisation and Writing: ODC

## 10 Acknowledgments

I would like to thank Francis-André Wollman for critical reading of the manuscript, as well as Richard Kuras and Yves Choquet for helpful discussions and advice. Katia Wostrikoff kindly provided plasmid CrRaf1-pJHL from which the G-rich linker sequence was amplified. Thank you also to Stéphane Lemaire and Pierre Crozet for suggesting that I should try PSAD-cTP.

